# Embryo-restricted responses to maternal IL-17A promote neurodevelopmental disorders in mouse offspring

**DOI:** 10.1101/2023.12.25.573296

**Authors:** David Andruszewski, David C. Uhlfelder, Genni Desiato, Tommy Regen, Carsten Schelmbauer, Michaela Blanfeld, Lena Scherer, Konstantin Radyushkin, Davide Pozzi, Ari Waisman, Ilgiz A. Mufazalov

## Abstract

Prenatal imprinting to interleukin 17A (IL-17A) triggers behavioral disorders in offspring. However, reported models of maternal immune activation utilizing immunostimulants, lack specificity to elucidate the anatomical compartments of IL-17A’s action and the distinct behavioral disturbances it causes. By combining transgenic IL-17A overexpression with maternal deficiency in its receptor, we established a novel model of prenatal imprinting to maternal IL-17A (acronym: PRIMA-17 model). This model allowed us to study prenatal imprinting established exclusively through embryo-restricted IL-17A responses. We demonstrated IL-17A transfer across the placental barrier and subsequent development of selected behavioral deficits in mouse offspring. More specifically, embryonic responses to IL-17A resulted in communicative impairment in early-life measured by reduced numbers of nest retrieval calls. In adulthood, IL-17A-imprinted offspring displayed an increase in anxiety-like behavior. We advocate our PRIMA-17 model as a useful tool to study neurological deficits in mice.

## Introduction

Maternal immune activation (MIA) during pregnancy is a recognized risk factor for neurodevelopmental disorders (NDDs) in offspring^1,2,3^. NDDs require lifelong and resource-intensive specialized care with no cure up to date^4^. The prevalence of NDDs is increasing, underlining the importance of developing safe treatment strategies suitable for use during pregnancy to prevent NDDs in the first place^5^. Preclinical studies demonstrated that MIA, induced through treatment of the pregnant female mouse with the synthetic viral RNA analog Poly(I:C), results in imprinting of offspring to develop NDDs. In this model, interleukin-17A (IL-17A), a cytokine produced by a subset of maternal T helper 17 (Th17) cells, emerged as a central player in the impaired neurodevelopment^6,7^. Despite a prominent role of IL-17A in Poly(I:C)-MIA, this model does not allow to study the direct effects of IL-17A on embryonic development due to broad immune activation, induction of additional maternal-derived mediators beyond IL-17A (such as IL-6, IL-1β, among others), pregnancy complications, and batch-to-batch Poly(I:C) variability^7,8^. More importantly, discerning whether effects on the offspring stem from embryonic or maternal responses to Poly(I:C)-induced IL-17A is challenging. This clarification is pivotal for devising therapies, given the potential risk posed to maternal and fetal health by unintended drug effects.

## Methods

Mice and colony maintenance All mice were on C57BL/6 background. Wild-type (WT) mice were bred in-house. IL-17RA knockout (IL-17RA^KO/KO^) mice were generated by crossing IL-17RA floxed mice^9^ with CMV-Cre mice^10^. Next, mice were intercrossed with IL-17A inducible (IL-17A^ind^) mice^11^ to generate IL-17RA^KO/KO^IL-17A^ind/WT^CMV-Cre^+/WT^ mice. These mice achieved constitutive germline IL-17A overexpression (IL-17A^OE^) on IL-17RA^KO/KO^ background. In the following intercrosses CMV-Cre was crossed out resulting in IL-17RA^KO/KO^IL-17A^OE/WT^ mice. To generate mice for control and experimental matings (Fig.1a), as well as for colony maintenance, IL-17RA^KO/KO^IL-17A^OE/WT^ mice were bred with IL-17RA^KO/KO^ mice.

**Fig. 1.**
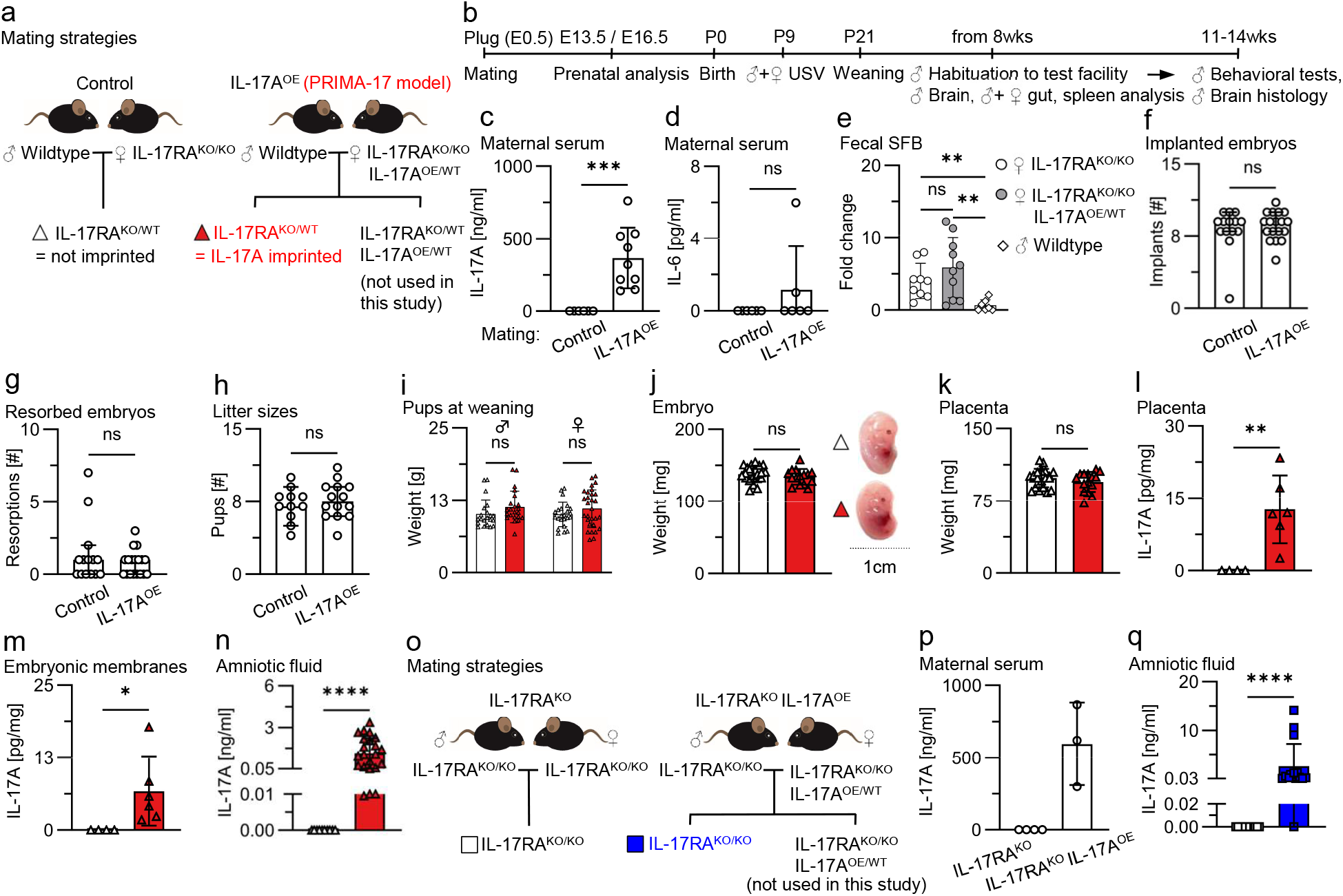
Analysis of a genetic model of maternal IL-17A overexpression combined with IL-17RA deficiency for pregnancy progression and IL-17A passage through the placenta. **a)** Mating scheme to achieve prenatal imprinting to maternal IL-17A (PRIMA-17 model). **b)** Experimental timeline of the study. Dams were sacrificed for prenatal analysis or allowed for natural delivery. At postnatal day 9 (P9), pups were subjected to an ultrasonic vocalization test (USV). At P21, pups were weaned and subjected to immune system analysis of the gut and spleen and brain analysis by flow cytometry in adulthood. A cohort of male offspring, from 8wks of age, were single-housed, transferred to the behavioral test facility for habituation and further subjected to behavioral tests and histological examination of the brain. **c)** Maternal IL-17A serum concentrations at E13.5. n=6 dams in control mating and n=9 in IL-17A^OE^ mating. **d)** Maternal IL-6 serum concentrations at E13.5. n=6 dams per group. **e)** Fecal SFB abundance in indicated breeders. n=9 (female IL-17RA^KO/KO^), n=10 (female IL-17RA^KO/KO^IL-17A^OE/WT^), n=7 (male wildtype). **=p<0.01, ns=p>0.05 Brown-Forsythe and Welch ANOVA with Dunnett’s T3 multiple comparisons test. **f)+g)** Number of implanted and resorbed embryos in indicated mating types. Data pooled from timepoints E13.5 and E16.5. n=14 litters in control mating and 18 litter in IL-17A^OE^ mating. **h)** Number of offspring born in indicated mating types. n=11 litters in control and n=14 litters in IL-17A^OE^ mating. **i)** Body weight of pups at weaning. n=21 (males) and 23 (females) from 7 litters in control mating (white), and 22 (males) and 29 (females) from 12 litters in IL-17A^OE^ mating (red). ns=p>0.05 two-way ANOVA with Šídák’s multiple comparisons test. **j)+k)** Body and placenta weight of embryos at E13.5. n=21 from 3 litter in control mating (white) and n=16 from 5 litters in IL-17A^OE^ mating (red). **l)+m)** IL-17A levels in placental and embryonic membrane (pooled umbilical cord and amniotic sac) tissue lysate at E13.5. n=4 from 1 litter in control mating (white) and 6 from 2 litters in IL-17A^OE^ mating (red). **n)** IL-17A levels in the amniotic fluid at E13.5. n=10 from 5 litters in control mating (white) and 24 from 13 litters in IL-17A^OE^ mating (red). **o)** Mating schemes to test maternal IL-17A passage through the placental barrier in IL-17RA deficient dams. **p)** Maternal IL-17A serum concentrations at E13.5. n=4 dams in IL-17RA^KO^ mating and 3 in IL-17RA^KO^ IL-17A^OE^ mating. **q)** IL-17A levels in the amniotic fluid at E13.5. n=12 from 4 litters in IL-17RA^KO^ mating (white) and 15 from 3 litters in IL-17RA^KO^ IL-17A^OE^ mating (blue). c)-h)+p) Data points represent individual mice. i)-n)+q) Data points represent individual pups or embryos. f)-h) Bar graphs indicate median±95% CI. c)-e)+i)-n)+p)-q) Bar graphs indicate mean±SD. c)+d),l)-n) ****=p<0.0001, ***=p<0.001, **=p<0.01, *=p<0.05, ns=p>0.05 two-tailed unpaired t-test with Welch’s correction. g)-h)+q) ****=p<0.0001, ns=p>0.05 two-tailed unpaired Mann-Whitney test. j)+k) ns=p>0.05 two-tailed unpaired t-test.

All mice were bred and maintained under specific-pathogen-free (SPF) conditions with food and water ad libitum in a 12 h light/12 h dark cycle with controlled room temperature and humidity at the Translational Animal Research Center (TARC) of the University Medical Center Mainz. After weaning, sex-matched littermates were co-housed until further use. To control for environmental factors, all breedings were conducted in the same area of the animal facility. Ethical approval was obtained from the Landesuntersuchungsamt Koblenz (G22-1-003), and all animal experiments followed the guidelines of the German Animal Welfare Act.

### Genotyping

Tissue biopsies were incubated in lysis buffer (Tris-HCl pH=8, KCl, EDTA, Igepal, Tween20 in ddH_2_O) supplemented with Proteinase K (7BioScience) at 56 °C on a shaker overnight. The digested tissue was incubated for 5 min at 95 °C, followed by centrifugation (12000 rcf, 10 min, RT). The resulting supernatant was used for genotyping through REDTaq PCR Reaction Mix (Sigma Aldrich). Specific primer sets were applied to detect the CMV-Cre and IL-17A transgenes, IL-17RA^WT^ and IL-17RA^KO^ alleles, and to determine the configuration of the Gt(ROSA)26Sor locus targeted with the IL-17A transgene.

### Timed pregnancies and litter size evaluation

One or two virgin females (8-10 weeks) were paired with one wild-type male in the evening (6-9 pm). The presence of a vaginal plug in the morning (7-10 am) indicated copulation and marked gestational day (E) 0.5. Females with mating plugs were separated from males and were housed alone or with a companion female. Pregnant females at term were inspected in the mornings to evaluate litter sizes.

### Maternal serum collection and prenatal analysis

Pregnant mice were sacrificed at E13.5 or E16.5 through deep anesthesia followed by cardiac blood withdrawal. Blood was incubated at room temperature for 30 min and centrifuged (12000 rcf, 20 min, RT) for serum collection. Sera were stored at -80 °C until used for IL-17A measurements. Uteri were extracted and washed with DPBS (Gibco) for assessment of embryo implantations and resorptions.

Fecal bacterial DNA isolation and segmented filamentous bacteria (SFB) quantification Prior to mating setup, fresh stool samples were collected, snap frozen in liquid nitrogen, and stored in -80 °C. A QIAamp® Fast DNA Stool Mini Kit (Qiagen) was used to isolate bacterial DNA. Then, by using a GoTaq® qPCR Master Mix (Promega) with 30 ng sample DNA, the relative amount of universal bacterial (primer: 5’-ATT ACC GCG GCT GCT GGC-3’ and 5’-ACT CCT ACG GGA GGC AGC AGT-3’) and SFB-specific (primer: 5’-GAC GCT GAG GCA TGA GAG CAT-3’ and 5’-GAC GGC ACG GAT TGT TAT TCA-3’) 16S rDNA was quantified. The fold change of SFB-specific DNA within the sample was calculated in reference to the universal bacterial DNA.

### Embryonic tissue collection

Individual embryos with their amniotic sac intact were dissected, washed with DPBS, and positioned on clean petri dishes. Amniotic fluid was collected by puncturing the amniotic sac with single-use 30 G insulin syringes (BD Biosciences), followed by subsequent centrifugation (300 rcf, 7 min, 4 °C) and collection of the cell-free supernatant. Placentas and embryonic membranes (pooled umbilical cord and amniotic sac) were separated from the embryo. Embryos, placentas, and membranes were dried on paper towels before being weighted. Deciduas were mechanically removed from placentas. The collected amniotic fluid and extra-embryonic tissues were stored at -80 °C until used for IL-17A measurements.

### Tissue processing and ELISA

Extra-embryonic tissues were lysed in RIPA buffer (Tris-HCl pH=7.5, NaCl, EDTA, Igepal, Sodium Deoxycholate in ddH_2_O) supplemented with protease-(cOmplete^TM^, Sigma-Aldritch) and phosphatase-(PhosSTOP^TM^, Roche) inhibitors using bead tubes (MP Biomedicals). After homogenization and centrifugation (12000 rcf, 10 min, 4 °C), the debris-free supernatant was collected. Cytokine concentrations in sera, amniotic fluid, and tissue homogenates were measured using an IL-6 (BD #555240) or IL-17A ELISA kit (Invitrogen #88-7371) according to the manufacturers recommendations. Concentrations were calculated with standard curves and final concentrations were normalized to tissue weight. Concentrations for background subtracted OD values below 0 were defined as 0 pg/ml.

### Ultrasonic vocalization test with pups to assess early social communication deficits

The day of birth was considered as postnatal day 0 (P0). On P9, breeding cages containing dam with litter were transferred to the testing room and habituated for 30 min. Before measurement of an individual litter, a beaker was cleaned once with 70% EtOH and dried using paper towels. After a 5 min evaporation period, a clean paper towel was placed in the beaker. One pup at a time was individually separated from the nest and placed into the beaker on top of the clean paper towel within a light-proof Styrofoam box. Vocalization profiles within the 0-125 kHz spectrum were recorded for 3 min using a Avisoft Bioacoustic UltraSoundGate 116Hb recorder with condenser microphone or a Metris Sonotrack control unit with microphone. After measurement, the pup was returned to its nest. After each pup of a litter, the paper towel was removed, and a fresh paper towel was placed inside the jar.

### Behavioral tests with adult mice

Tests with adult mice were conducted and analyzed at the Mouse Behavior Unit (MBU) of the Leibniz Institute for Resilience Research (LIR) gGmbH, Mainz, Germany. Male mice at 8-10 weeks were transferred in separated cages for single-housing in the testing facility. There, mice acclimated for at least one week to the new environment. During this period, mice were accustomed to the experimenter through repeated handling. For each test, mice were introduced to the testing room and acclimated for 30 min before being gently transferred to the test area (RT, 70 Lux, 9 am - 6 pm) via tunnel or cup handling. Except for the marble burying test, all tests were video-recorded and subjected to automated analysis. The testing areas were cleaned between individual mice. At least one day of rest separated consecutive tests. The sequence of tests followed the description below.

#### Elevated plus maze test to assess anxiety-like behavior

A mouse was placed at the intersection point of an elevated X-shaped maze (75 cm above the ground). It consisted of two arms (77 cm x 7 cm), one with walls (20 cm high, closed arm), and the other without (open arm). The mouse explored both arms for 10 min, and time spent and number of entries into each arm were quantified.

#### Open field test to assess anxiety-like behavior and locomotor activity levels

A mouse was introduced into an empty test arena (42 cm x 42 cm x 41 cm) and allowed to explore for 10 min. The time spent in a defined center area (10 cm x 10 cm) was quantified.

#### Marble burying test to assess repetitive and anxiety-like behavior

A mouse was placed in a test area (22 cm x 38 cm x 20 cm) filled to a depth of 5 cm with bedding. On top of the bedding, 16 marbles were placed in a specific pattern (4 marbles per row, 4 rows). The test mouse was free to explore the cage and dig in the bedding. After 30 min, the mouse was removed, and the number of fully buried marbles was counted.

#### Three-chamber sociability and social novelty test to assess social interaction deficits

After a 10 min habituation in an empty test arena (63 cm x 41 cm x 22 cm) with three chambers (21 cm x 41 cm x 22 cm each), the test mouse was placed in the central chamber. Additionally, an unfamiliar adult wild-type male mouse was introduced in one of the side chambers, while an inanimate object was placed in the other. During a 10 min period, the test mouse interacted with both the mouse and object. Interaction time with the mouse and object was quantified. This process was repeated with a novel (unfamiliar) male mouse replacing the object. Interaction time with familiar and novel mice was quantified. Unfamiliar mice were used for up to three consecutive encounter with experimental mice.

### Leukocyte isolation from intestinal lamina propria and spleen

Cells were isolated as described in full detail elsewhere^12^. In brief: Colons and small intestines (without caecum) from male (9 weeks) or female mice (8-20 weeks) were isolated and transferred into RPMI-1640 (Gibco) medium supplemented with L-glutamine, Penicillin/Streptomycin (Pen/Strep), FCS and HEPES (hereafter referred to as 3% medium). Peyer’s patches were removed, and intestines were cleaned with 3% medium before incubation (37 °C, 20 min, horizontal shaker) with strip medium (3% medium supplemented with EDTA and DTT). Samples were vortexed rigorously and the strip medium was removed by transferring the samples on a strainer. The tissue was transferred into RPMI-1640 medium supplemented with L-glutamine, Pen/Strep, EDTA, and HEPES and vortexed rigorously. The medium was removed by transferring the samples on a strainer and the tissue washed with 3% medium. Then intestines were mechanically dissociated and incubated (37 °C, 30 min, horizontal shaker) in digestion medium (RPMI-1640 supplemented with L-glutamine, Pen/Strep, β-ME, HEPES, Sodium Pyruvate, Liberase TL^TM^ (Sigma Aldritch) and DNase I). Then, the cell suspension was vortexed and passed through a 100 μm filter. Finally, cells were centrifuged (450 rcf, 10 min, 4 °C), resuspended in 3% media and kept on ice until further use.

Spleens were mechanically dissociated through a 40 μm filter and resuspended in DPBS with 2% v/v FCS (hereafter referred to as PBS/FCS). Cells were centrifuged (300 rcf, 7 min, 4 °C) and resuspended in 1 ml Ammonium-chloride-potassium lysis buffer to remove erythrocytes. After 1 min incubation at RT, cells were washed, centrifuged (300 rcf, 7 min, 4 °C) and resuspended in PBS/FCS.

### Microglia isolation from brains

Cells were isolated as described in full detail elsewhere^13^. In brief: deeply anesthetized male mice (7-8 weeks) were transcardially perfused (0.9% w/v NaCl), decapitated, and their brain was removed from the skull. Brains were mechanically dissociated with a razor blade and resuspended in digestion solution containing 1.5 mg/ml Collagenase II (Gibco) and 50 μg/ml DNase I (Roche) in PBS with Calcium and Magnesium (Gibco). After incubation (30 min, 37 °C) in water bath, the reaction was stopped, the lysates were homogenized and centrifugated (300 rcf, 5 min, 4 °C), and the supernatant aspirated. Then, the pellets were resuspended in 70% Percoll (Cytiva) and a three layer 30%/37%/70% Percoll gradient separation was performed by centrifugation at 40 min, 500 rcf, RT. Consequently, the upper myelin-enriched layer was removed, and leukocytes were harvested from the interphase of the middle 37% and lower 70% Percoll fractions. Cell suspensions were centrifugated (300 rcf, 5 min, 4 °C) and the pellets resuspended in PBS/FCS.

### Flow cytometry

Cells were centrifuged (300 rcf, 7 min, 4 °C), resuspended in PBS/FCS supplemented with Fc-block (BioXCell) and incubated for 15 min on ice. After, cells were centrifuged (300 rcf, 7 min, 4 °C), resuspended in PBS/FCS containing fixable viability dye (APC-eFl780, eBioscience #65-0865, 1:1000) and antibodies directed against surface molecules CD11b (biotinylated, eBioscience #13-0112, 1:400 or PE-Cy7, BioLegend # 101216, 1:1000), CD45 (BV510, BioLegend #103138, 1:300), F4/80 (PE, eBioscience #12-4801, 1:50), CD4 (BV510, Biolegend #100559, 1:200), TCRβ (FITC, Biolegend #109205, 1:200), MHCII (FITC, BioLegend #107605, 1:1000), CD86 (biotinylated, BD Biosciences #553690, 1:600). After 15 min incubation on ice, cells were centrifuged (300 rcf, 7 min, 4 °C) and fixed (1 h, 4 °C) using the eBioscience Foxp3/Transcription factor staining buffer set (ThermoFisher #00-5523-00) fixation/permeabilization reagent or the Cytofix/Cytoperm Fixation/Permeabilization Kit (BD Biosciences #554714) Fix/Perm solution. After incubation, cells were centrifuged (350 rcf, 7 min, 4 °C) and resuspended in eBioscience or Cytofix/Cytoperm permeabilization buffer containing Streptavidin (PerCP, BioLegend #405213, 1:600) and/or antibodies directed against intracellular targets Foxp3 (APC, eBioscience #17-5773, 1:200), RORγt (BV421, BD Biosciences #562894, 1:200), CD68 (APC, BioLegend #137007, 1:400), IgG2a isotype control (APC, eBioscience # 17-4321, 1:400), and Ki67 (APC, BioLegend #646407/8, 1:400). After 1 h incubation at 4 °C, cells were centrifuged (350 rcf, 7 min, 4 °C), resuspended in PBS/FCS and acquired on a BD FACSCanto II flow cytometer.

### Brain histology

After undergoing behavioral tests, male mice were deeply anesthetized and transcardially perfused with 0.9% w/v NaCl followed by 4% PFA (Santa Cruz). Perfused mice were decapitated, and their skull was post-fixed and stored in 4% PFA. Then, brains were removed from the skull and coronal 50μm -thick slices were cut with a VT1000S vibratome (Leica Microsystems). After cutting, slices were washed with cold PBS followed by permeabilization and nonspecific binding sites blockage with 1% BSA w/v, and 0.3% Triton-X v/v in PBS for 90 min at RT. Afterwards, sections were incubated with primary rabbit anti-IBA1 antibody (WAKO Chemicals #019-19741, 1:500), 1% BSA, and 0.1% Triton-X in PBS overnight at 4 °C. Slices were then rinsed 3 times with PBS and incubated with Alexa Fluor-488 conjugated anti-rabbit secondary antibodies (Thermo Fisher Scientific # A-11008, 1:500), 1% BSA, and 0.1% Triton-X in PBS, for 2 h at RT protected from light. Slices were then washed, counterstained with Hoechst-33342 (Thermo-Fisher #62249, 1:1000) and mounted with Fluorsave mounting medium (Millipore). Confocal images were acquired using a Leica SP8II laser scanning confocal microscope equipped with an HC PL APO 20x/0.75 CS2 objective and ACS APO 40x/1.40 oil immersion objective. For microglia density quantification, at least four z-stack acquisitions per hemisphere and animal of the primary somatosensory area (S1) and at least two sections for each animal were considered. IBA1 positive cells were counted, and the density was quantified by normalizing the number of microglia to the acquired volume. The analyzed regions of one animal were averaged and presented as single data point.

### Statistical analysis and data presentation

Discrete datasets and non-normal distributed continuous datasets (normal distribution was tested in conspicuous datasets via Kolmogorov-Smirnov test) were analyzed using the two-tailed Mann-Whitney test to determine statistical significance. Continuous datasets with a presumed normal distribution underwent analysis via the two-tailed unpaired t-test. Welch’s correction was applied when significant differences in variance, tested via F test, were present between two groups. For the statistical comparison of more than 2 groups and one variable, one-way ANOVA test with correction for multiple comparisons was used. For datasets with more than two groups and more than one variable, two-way ANOVA with correction for multiple comparisons was used. No randomization and no blinding were used throughout the study.

Data are presented either as median±95% confidence interval (CI) or mean ± standard deviation (SD), depending on whether the data were discrete or continuous. Statistical significance levels were indicated as follows: ns=p>0.05, *=p<0.05, **=p<0.01, ***=p<0.001 and ****=p<0.0001. Behavioral test results involving adult mice were presented using box and whisker plots, encompassing 2.5-97.5 percentiles and min/max values.

Group sizes were determined based on various criteria. For cytokine measurements, at least three mice/embryos per group were included, except for amniotic fluid IL-17A, which required at least 10 embryos from three independent litters. Prenatal analysis, fecal SFB, litter sizes, pup weight at weaning, and ultrasonic vocalization analysis utilized all available mice within specific timeframes. Behavioral tests in adults were sample-sized using an a priori power analysis (α = 0.05, β = 0.1, Cohen’s d = 1). Flow cytometry and histological analyses included at least 3 and 4 animals per group, respectively, based on experimental experience and availability of mice. Technical replicates were averaged and presented as a single datapoint, thus, each datapoint represents an individual mouse/embryo. The exact sample size (n) for each dataset can be found in the figure legends

### Software

Avisoft SAS Lab Pro software 5.1 or Metris Sonotrack 2.7.0 was used for ultrasonic vocalization profile recording. Metris Sonotrack 2.7.0 was used for ultrasonic vocalization profile analysis. G*Power v3.1.9.7 was used for determining group sizes for behavioral tests. EthoVision XT by Noldus was used for automated analysis of behavioral test video recordings. Graphs were generated, and statistical analyses were performed using GraphPad Prism 9. Illustrations were created using CorelDRAW Graphics Suite 2022. Microsoft 365 (Office) tools, specifically Word for manuscript writing, Excel for data analysis, and Publisher/PowerPoint for figure design, were used.

## Results

The aim of our study was to establish a novel model of prenatal imprinting to maternal IL-17A (acronym: PRIMA-17 model) and use it to investigate whether embryo-restricted responses to IL-17A lead to behavioral disturbances. To achieve this, we defined the following criteria for the model: 1) maternal blood should be enriched for IL-17A; 2) embryonic tissues should be responsive to IL-17A; 3) maternal tissues should be unresponsive to IL-17A; 4) pregnancy outcome and nursing should remain unaffected. To meet these criteria, we employed a genetic approach that minimized the need for mouse handling, compared to pharmacological interventions, thereby reducing maternal stress during pregnancy. For that, we generated mice with constitutive germline IL-17A overexpression (OE) from a single copy transgene inserted into the ROSA26 locus while the second ROSA26 allele remained wild-type (WT). Additionally, we converted the IL-17 receptor A (IL-17RA) in these mice to the complete knockout (KO) state (referred to as IL-17RA^KO/KO^IL-17A^OE/WT^ mice). Deficiency in IL-17RA in the mother ensures maternal unresponsiveness to IL-17A^14^ and therefore, IL-17A overexpression should not cause any maternal stress during pregnancy and nursing. Subsequently, IL-17RA^KO/KO^IL-17A^OE/WT^ females were paired with WT males (termed IL-17A^OE^ experimental mating) to produce offspring with two distinct genotypes: IL-17RA^KO/WT^IL-17A^WT/WT^ (short IL-17RA^KO/WT^) and IL-17RA^KO/WT^IL-17A^OE/WT^, both responsive to IL-17A due to one functional IL-17RA wild-type allele inherited from the father (Fig.1a). Transgenic IL-17A overexpression on IL-17RA wild-type background leads to moderate-to-lethal inflammation^11,15^ and therefore IL-17A transgenic IL-17RA^KO/WT^IL-17A^OE/WT^ offspring were not considered for the purpose of the present study. Instead, we evaluated IL-17RA^KO/WT^ littermate offspring without IL-17A transgene which were exposed to maternal-derived IL-17A *in utero*, allowing to unequivocally study the direct role of prenatal imprinting to IL-17A. To generate appropriate controls for IL-17A-imprinted offspring, IL-17RA^KO/KO^ females without IL-17A transgene were paired with WT males (termed control mating). This mating yielded genotype-matched IL-17RA^KO/WT^ offspring that developed in the absence of transgenic IL-17A in dams (Fig.1a). Consequently, we performed a set of pre- and postnatal analyses at different time points to determine the effect of embryonic IL-17A responses on pregnancy outcome, immune system homeostasis, behavior and histological analysis of the brain (Fig.1b).

As expected, serum of IL-17A transgenic dams in experimental IL-17A^OE^ matings displayed high levels of IL-17A (Fig.1c). Importantly, no such elevation was observed in the levels of IL-6 (Fig.1d), yet another critical cytokine for NDD development^6,7^. A disrupted gut microbiota homeostasis in dams promotes behavioral abnormalities in offspring exposed to MIA^6,16,17^. To consider the potential impact of the maternal microbiota on prenatal imprinting in the PRIMA-17 model, we used IL-17RA^KO/KO^ and IL-17RA^KO/KO^IL-17A^OE/WT^ dams which were littermates and were continuously co-housed prior to mating set up. Co-housing is a useful strategy to ensure aligned microbiota composition in mice^18^. To confirm the alignment of the maternal microbiota, we quantified the amount of segmented filamentous bacteria (SFB), a commensal bacterium that is essential for the development of behavioral disorders in the Poly(I:C) MIA model^6^. For that we collected feces from the breeding partners one day before mating set up, isolated bacterial DNA and performed qPCR with primers specific to SFB. We found no differences in the abundance of SFB between IL-17RA^KO/KO^ and IL-17RA^KO/KO^IL-17A^OE/WT^ breeder females (Fig.1e). Of note, IL-17RA deficiency in females resulted in higher SFB abundance compared to their wild-type male mating partners, which is in line with our previous report describing an increase in SFB in IL-17A deficient mice^18^. We concluded that our mating set up allows us to rule out potential effects related to differences in maternal gut microbiota and attribute findings solely to embryonic responses to maternal IL-17A.

Since complications during pregnancy can lead to behavioral disorders in offspring^19,20^, we evaluated pregnancy progression and outcome in our model. Our analysis revealed that high levels of IL-17A in dams devoid of IL-17A signaling does not affect the number of implanted and resorbed embryos (Fig.1f-g). Litter sizes at wean remained similar in both types of matings, indicating that neonatal survival of pups born to IL-17RA^KO/KO^IL-17A^OE/WT^ dams remained normal (Fig.1h). The body weight of IL-17A-imprinted pups at wean did not differ from controls (Fig.1i). Together these data imply that maternal nursing in the experimental IL-17A^OE^ mating remained unaffected. Furthermore, IL-17RA^KO/WT^ embryos generated in IL-17A^OE^ matings exhibited unaltered gross morphology, body- and placental weight at E13.5 (Fig.1j-k). We concluded that embryonic IL-17A responses in IL-17RA deficient dams do not result in adverse pregnancy outcomes.

In humans and mice, the placenta strictly separates maternal and embryonic blood^21,22^. To test whether maternal-derived IL-17A crosses the placenta, we measured IL-17A levels within embryonic tissues and amniotic fluid. We revealed elevated IL-17A levels in placentas, embryonic membranes, and amniotic fluid of IL-17RA^KO/WT^ embryos in IL-17A^OE^ matings, as compared to controls (Fig.1l-n). Because IL-17A ELISA cannot discriminate between transgenic and endogenous cytokine, IL-17A in the amniotic fluid could be of maternal origin, of fetal origin induced by maternal IL-17A, or both. To clarify the origin of IL-17A we modified the mating strategy by replacing wild-type males with IL-17RA^KO/KO^ males (Fig.1o). Offspring born in this mating scheme are completely unresponsive to IL-17A due to homozygous IL-17RA deficiency. As a result, maternal-derived IL-17A cannot induce the expression of embryonic IL-17A. First, we confirmed that IL-17RA^KO/KO^IL-17A^OE/WT^ dams used in this experimental setup indeed express high levels of IL-17A (Fig.1p). Second, we analyzed amniotic fluid and detected elevated levels of IL-17A in the amniotic fluid of IL-17RA^KO/KO^ embryos recovered from IL-17RA^KO/KO^IL-17A^OE/WT^ dams at E13.5 (Fig.1q). Due to complete deficiency in IL-17RA and inability of IL-17A to be induced on site, this data confirms that maternal IL-17A can penetrate the placental barrier in our system. Together, we concluded that elevated IL-17A which we observed in the original IL-17A^OE^ mating (Fig.1a, n) is, at least in part, of maternal origin and will serve as a driving force for prenatal imprinting in the PRIMA-17 model.

Prenatal imprinting resulting from maternal infection can alter immune responses in the gut of offspring^23,24^. This way, maternal-derived IL-6 can act on embryonic intestinal stem cells, thereby leading to persistent changes in the Th17 cell population in adulthood^24^. To explore a potential effect of maternal-derived IL-17A on T cell homeostasis in our PRIMA-17 model, we assessed RORγt^+^ Th17 cells, Foxp3^+^ regulatory T (Treg) cells as well as gut-associated RORγt^+^Foxp3^+^ Treg cells in IL-17RA^KO/WT^ offspring born in control and IL-17A^OE^ matings. We analyzed male and female mice separately and observed no variations in the distribution or quantity of these T cell subsets, neither in the small intestine, nor in the colon or spleen (Fig.2 and Supplementary Fig.1). Taken together, we found no indication for imprinting effects of maternal IL-17A on the intestinal and systemic Th17/Treg cell homeostasis.

**Fig. 2.**
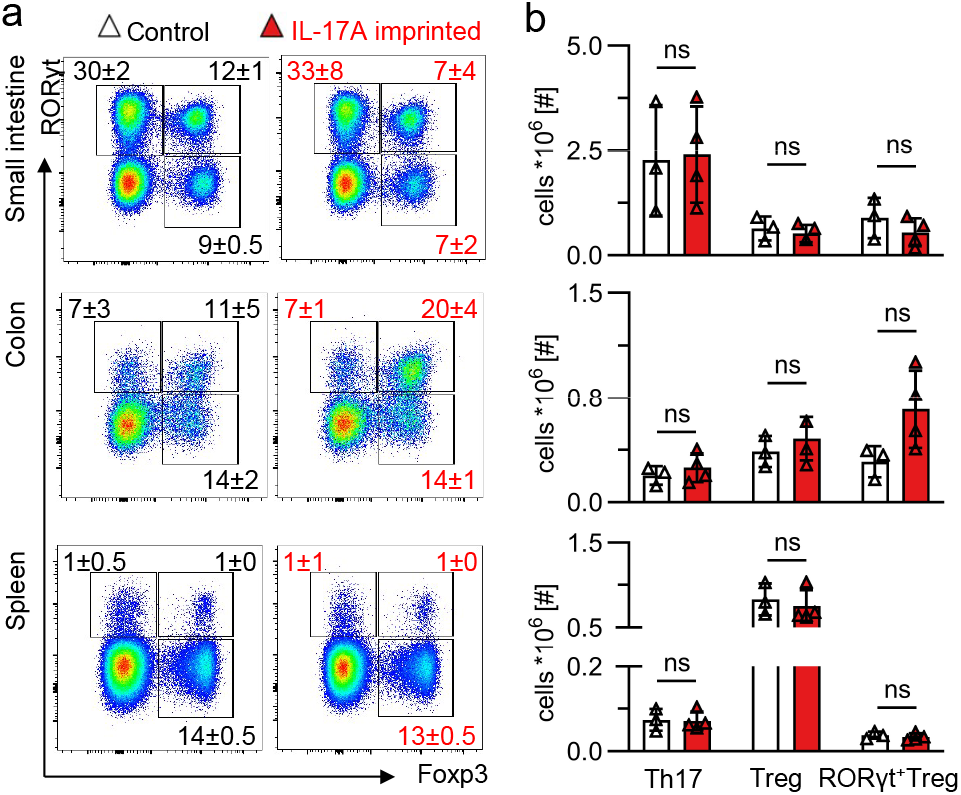
Th17 and Treg cells are not impacted in adult offspring prenatally exposed to IL-17A. **a)** Representative FACS plots depict RORγt and Foxp3 expression by T cells (gated on: TCRβ^+^CD4^+^ cells) in male mice. Numbers within plots indicate mean±SD frequencies of cells in the adjacent gate. No statistically significant differences were noted in frequencies between groups. **b)** numbers of Th17, Treg and RORγt^+^ Treg cells in the small intestine (top), colon (middle) and spleen (bottom). For each organ, n=3 from 1 litter born in control mating (white) and n=4 from 2 litters born in IL-17A^OE^ mating (red). Datapoints represent individual mice. Bar graphs indicate mean±SD. ns=p>0.05 two-tailed unpaired t-test. Control=IL-17RA^KO/WT^ offspring born to IL-17RA^KO/KO^ dams (white). IL-17A imprinted=IL-17RA^KO/WT^ offspring born to IL-17RA^KO/KO^IL-17A^OE/WT^ dams (red).

Next, we assessed whether embryo-restricted IL-17A signaling impacts behavior of offspring. Therefore, we first evaluated the ultrasonic vocalization (USV) profile of pups at postnatal day 9 (P9). Vocalizations emitted by pups upon nest removal are considered as a type of social communication to gain the mother’s attention and achieve retrieval to the nest^25^. IL-17A-imprinted pups showed reduced number of vocalizations compared to control offspring (Fig.3a). At the same time, the duration of nest retrieval calls was not affected in pups born in IL-17A^OE^ mating (Fig.3a). A sex-specific analysis revealed a reduction in the number of calls in IL-17A imprinted pups of both gender. In contrast, the duration and frequency range were not changed in IL-17A imprinted pups, neither in males nor in females (Supplementary Fig.2). The communicative irregularities in early life led us to conduct behavioral tests with adult mice. For better comparison with reported findings, we focused our further analysis on male offspring as studies with MIA predominantly use males^6,7,26,27^. To assess anxiety-like behavior, we performed an elevated plus maze test and found no differences in the time spent in open or closed arms in male offspring born in IL-17A^OE^ matings (Fig.3b). However, the open-field test revealed a reduction in the time spent in the center area in IL-17A-imprinted mice (Fig.3c). This effect did not result from changes in locomotor activity as the distance mice traveled during the test remained similar to controls (Fig.3c). We concluded that prenatal exposure to IL-17A may lead to anxiety-like behavior under specific experimental settings. Next, we assessed sociability in adult mice. No differences were noted in the time spent with inanimate objects or unfamiliar mice during a social approach test. Similarly, a social novelty test did not reveal differences in the interaction time with familiar or novel mice (Fig.3d-e). These results pointed out that social approach behavior and social memory was not affected in IL-17A-imprinted offspring. Finally, a reduction in number of buried marbles was observed in the marble burying test, indicating a reduction in repetitive digging behavior (Fig.3f). Collectively, our data show that embryonic IL-17A signaling alone is sufficient to cause a selective set of behavioral disturbances, as was evident in the ultrasonic vocalization test, the open field test, and the marble burying test.

**Fig. 3.**
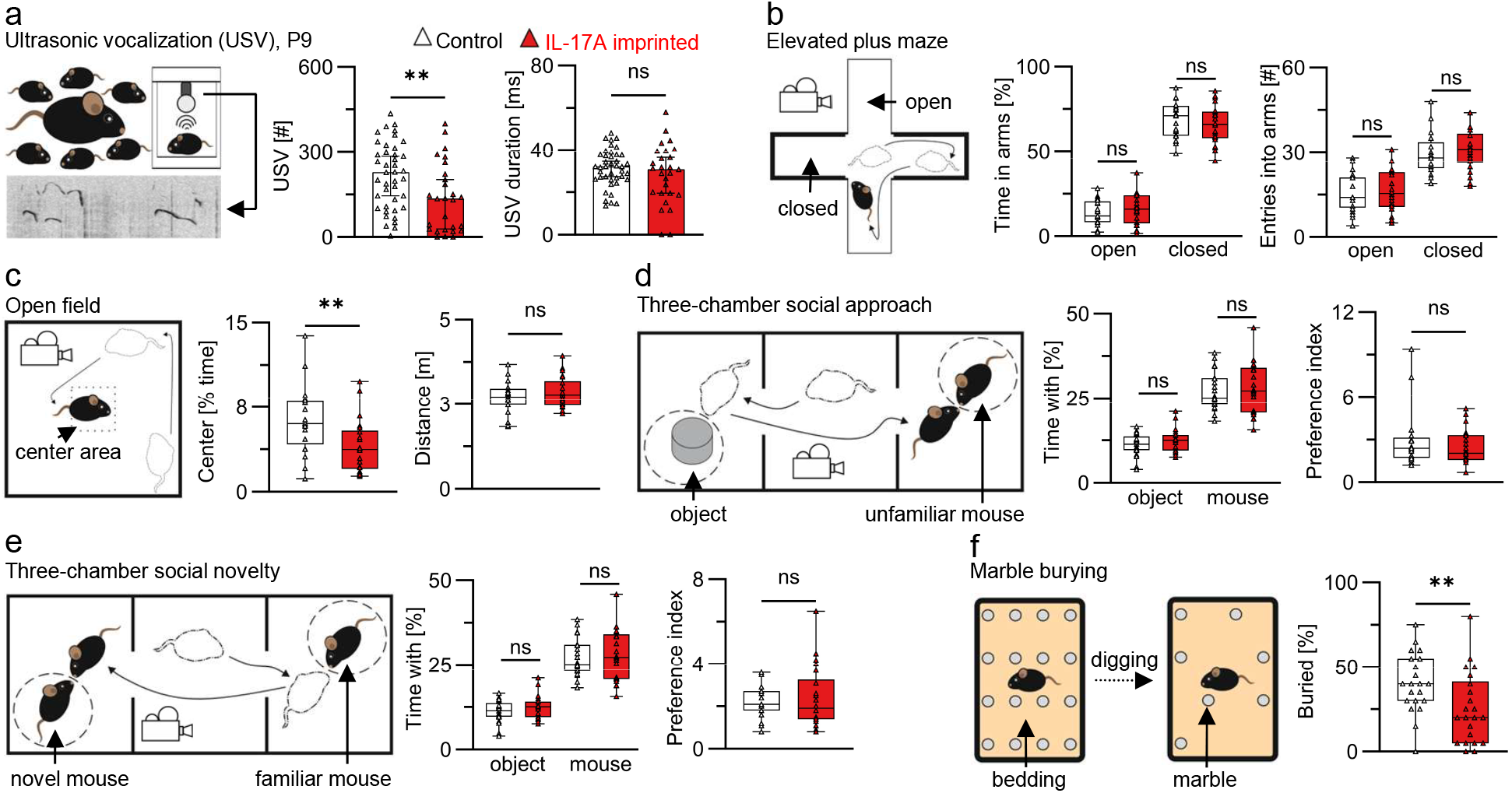
Embryo-restricted IL-17A responses lead to behavioral disturbances. **a)** Ultrasonic vocalization test. Left: Number of ultrasonic vocalizations (USV) emitted in the test period, median±95%CI. Right: Duration of vocalizations, mean±SD. Datapoints represent individual pups of mixed gender at P9. n=41 pups from 7 litters born in control (white) and 27 pups from 8 litters born in IL-17A^OE^ matings (red) **b)** Elevated plus maze test. Left: Time spent in open and closed arms. Right: Number of entries into open and closed arms. **c)** Open field test. Left: Time spent in the center area. Right: Total distance moved within the test period. **d)** Three-chamber social approach test. Left: Time spent with an inanimate object and an unfamiliar mouse. Right: Preference index as ratios of time spent with an unfamiliar mouse to time spent with an inanimate object. **e)** Three-chamber social novelty test. Left: Time spent with novel and familiar mice. Right: Preference index as ratios of time spent with an unfamiliar mouse to time spent with a familiar mouse. **f)** Marble burying test. Frequency of buried marbles within the test period. a)+f) **=p<0.01, ns=p>0.05 two-tailed unpaired Mann-Whitney test. b)-e) **=p<0.01, ns=p>0.05 two-tailed unpaired t-test. b)-f) Boxes indicate 2.5-97.5 percentile. Whiskers indicate Min./Max. values. Horizontal lines indicate median. n=21 from 7 litters (white) and 22 from 12 litters (red). Pooled data from 4 independent test cohorts. Males only, 11-14wks. Graphics on the left show the experimental setup of an individual test. Control=IL-17RA^KO/WT^ offspring born to IL-17RA^KO/KO^ dams (white). IL-17A imprinted=IL-17RA^KO/WT^ offspring born to IL-17RA^KO/KO^IL-17A^OE/WT^ dams (red). Data points in all graphs represent individual mice. Behavioral tests were performed in sequence as described in Materials and Methods.

Microglia is brain-resident immune cell type important for neuronal function and behavior. MIA is known to affect microglia in offspring^28,29^. To gain mechanistic insight on how maternal-derived IL-17A disrupts behavior in offspring we opted to analyze microglia in the brain of mice born to IL-17A^OE^ females in our PRIMA-17 model. First, we isolated microglia from the full brain and performed flow cytometry. We defined microglia as the CD45^+^CD11b^int^ cell population and found no differences in their frequency and numbers between IL-17A-imprinted and control mice (Fig.4a-c). In accordance with this observation, expression of the proliferation marker Ki67 remained unaltered in microglia (Fig.4d). Furthermore, we found no difference in the expression of activation markers MHCII, CD86 (Fig.4e-f), and a functional marker CD68 (Fig.4g-h) in microglia recovered from male offspring born in our PRIMA-17 model. The primary somatosensory cortex (S1) was previously identified as an affected region in the Poly(I:C)-MIA model^6,7,30^. Therefore, we performed a histological examination of IBA1^+^ microglia in the S1 region but failed to detect a difference in microglia density (Fig.4i). Collectively, this data implies that homeostatic microglia most likely is not affected by prenatal imprinting to IL-17A in the PRIMA-17 model and is unlikely to cause behavioral deficits in offspring born to IL-17A^OE^ females.

**Fig. 4.**
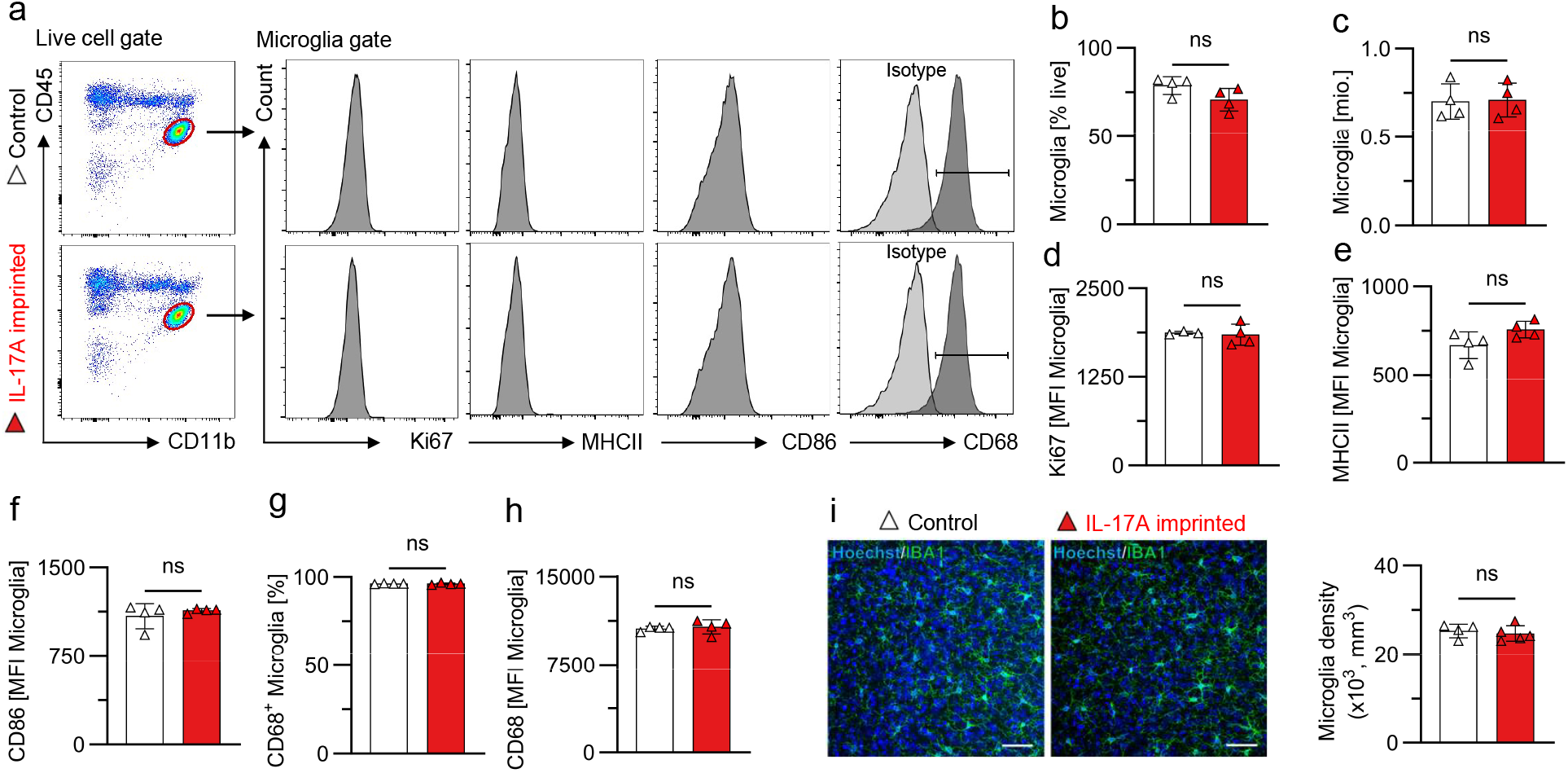
Microglia analysis of IL-17A-imprinted adult mice. **a)** Representative FACS plots and histograms of microglia analysis in IL-17A-imprinted and control mice. Histograms show the expression of indicated markers Ki67, MHCII, CD86 and CD68 (including isotype control and positive gate). **b)+c)** Microglia frequency among live cells and total microglia number obtained from the full brain. **d)** Mean fluorescent intensity (MFI) of Ki67 expression. **e)** MFI of MHCII expression. **f)** MFI of CD86 expression. **g)+h)** Frequency of CD68^+^ Microglia and MFI of CD68 expression. **i)** Left: Representative confocal images of the primary somatosensory region S1 labelled with antibodies against IBA1, identifying microglia (green) and counterstained with Hoechst 33435 (blue). Scale bars = 50 μm. Right: Quantification of IBA1^+^ microglia density in the S1 region. Bar graphs indicate individual mice, mean±SD. n=4 mice from 2 litters in control mating and 4 from 2 litters (a-h), n=5 from 4 litters (i) in IL-17A^OE^ mating. a)-h) ns=p>0.05 two-tailed unpaired t-test. i) ns=p>0.05, two-tailed unpaired Mann-Whitney test. Control=IL-17RA^KO/WT^ offspring born to IL-17RA^KO/KO^ dams (white). IL-17A imprinted=IL-17RA^KO/WT^ offspring born to IL-17RA^KO/KO^IL-17A^OE/WT^ dams (red).

## Discussion

Here we developed an innovative model of prenatal imprinting to maternal IL-17A, which we termed by its acronym: PRIMA-17. The model is based on a unique type of mating scheme which involves IL-17A transgenic females on IL-17RA deficient background and allows to study embryo-specific responses to a single cytokine - IL-17A. PRIMA-17 holds an advantage over other models which mimic maternal immune activation but lack specificity to the induced stress factor(s).

By analyzing our PRIMA-17 model, we found no indication that embryonic IL-17A responses cause pregnancy complications, which contrasts to growth retardation observed in offspring born to WT dams treated with IL-17A^31^. However, unlike in our study, WT dams were IL-17A responsive and experienced handling stress, which may explain differences in pregnancy outcomes. We showed that maternal IL-17A passes through the placental barrier in our model. Although we cannot exclude the contribution of embryonic IL-17A potentially induced by maternal IL-17A, we concluded that in either case, the placental passage of maternal IL-17A is likely a critical mechanism through which IL-17A imprints behavioral deficits (Supplementary Fig.3). Studies performed with human placentas demonstrated that the placental barrier is permeable for some inflammatory cytokines (IL-6), but not for others (TNFα, IL-1β)^32^. A recent study found that fluorophore conjugated IL-17A could be detected within the embryo shortly after injection into pregnant mice^33^. The authors hypothesized that IL-17RA expression by embryonic trophoblast cells is required for the passage across the placental barrier. However, we show that the passage can occur also independently from maternal and embryonic IL-17RA expression (Fig.1o-q).

Previous studies reported social behavior impairments following maternal Poly(I:C) treatment^6,7^. Intriguingly, maternal IL-17RA deficiency prevented Poly(I:C)-induced disturbances in social behavior^7^. Likewise, we found no alterations in social behavior in IL-17A-imprinted offspring born to IL-17A^OE^ dams with IL-17RA deficiency in our PRIMA-17 model. These reported findings, together with ours, suggest that sociability in offspring is shaped by maternal rather than embryonic responses to IL-17A. In contrast, embryonic responses alone are sufficient to drive anxious behavior during the open field test, which overlap between Poly(I:C)-MIA and our genetic model^6^. Beyond the open field test, which is the standard to assess anxious behavior, elevated plus maze test and marble burying test could also be used to assess anxiety in mice. In our study, IL-17A-imprinted mice exhibited no signs of anxiety-like behavior in the elevated plus maze test, but showed a reduction in buried marbles which could be interpreted as anxious behavior^34,35^. All together, we concluded that embryonic imprinting to IL-17A influences anxious behavior in a context-dependent manner.

Two opposing effects were observed in other behavioral tests between reported Poly(I:C)-MIA offspring and offspring born in our model. More specifically, Poly(I:C)-MIA offspring emitted more ultrasonic vocalizations than controls, while IL-17A-imprinted offspring from the PRIMA-17 model emitted fewer. Also, IL-17A-imprinted mice buried fewer marbles, while Poly(I:C)-MIA offspring bury more marbles ^6,7^. These contradictory results might stem from differences in IL-17A dosage and exposure timing. Poly(I:C) is typically injected at midgestation and leads to a mild and transient elevation of maternal IL-17A level^6,7^. In contrast, our PRIMA-17 model is characterized by chronically elevated and rather high systemic IL-17A levels which supposedly may act on embryos starting from earliest developmental stages until birth. Unfortunately, a direct comparison of IL-17A level in human inflammatory conditions with IL-17A level in rodent models is challenging due to limited availability of reliable human data. Additional factors induced by Poly(I:C), like IL-6, might have synergistic effects with IL-17A, which would result in distinct behavioral outcomes compared to our IL-17A-specific model, which is devoid of such factors.

We hypothesize that maternal-derived IL-17A ultimately reaches the embryonic brain to imprint behavioral deficits. However, the exact molecular mechanisms leading to these changes remain to be elucidated. Previous studies found that Poly(I:C)-MIA activates the integrated stress response and thereby disrupts proteostasis in embryonic brains in an IL-17A dependent manner^27^. Another study suggested that IL-17A promotes the production of maternal-derived kynurenine, a metabolite with neuroactive properties, which presumably crosses the placental barrier^36^. While the former might be a potential mechanism also in our PRIMA-17 model, the latter seems to require maternal kynurenine production stemming from a maternal response to IL-17A, which does not occur in our model due to maternal deficiency in IL-17RA.

The absence of immune system alterations in the gut and spleen, together with unaffected microglia activation in the brain of IL-17A-imprinted offspring suggests that neurons might be a primary target for maternal-derived IL-17A. Previous reports showed that IL-17RA deletion in neurons of adult mice, driven by Syn1-Cre, impacts behavior^26,37^. Another study utilizing Nestin-Cre-specific IL-17RA deficient mice showed that IL-17A acts on neuronal and glial cell precursors^30^, but mechanistic evidence for IL-17A signaling in embryonic brain neurons still remains limited. Follow up studies should clarify IL-17A responsive cell types in the embryonic brain critical for the development of behavioral abnormalities.

Taken together, our innovative mouse model of prenatal imprinting to maternal IL-17A, allows to study embryonic responses to a single stressor, IL-17A. Our findings suggest that targeting the embryonic IL-17A pathway during pregnancy may prevent some forms of neurodevelopmental disorders, such as abnormalities in vocalization, anxiety-like and repetitive behavior, which are hallmarks of autism spectrum disorders.

## Data availability

Data used for graphs, raw data and statistical analysis can be provided upon reasonable request.

## Acknowledgements

We thank Dr. Ulrich Schmitt and Mona Flachsel (both MBU LIR, Mainz) for technical assistance with behavioral testing. We thank Elena Zurkowski for assistance with flow cytometry experiments. Finally, we thank Dr. Khalad Karram and Dr. Margarita Tevosian for the fruitful discussions.

## Contributions

D.A. and I.A.M. designed experiments, performed timed pregnancies, tissue and fecal collections, and analyzed data. D.A. and M.B. performed ELISA and flow cytometry experiments. G.D. and D.P. performed histological IBA1 analysis. T.R. advised on microbiota analysis and isolated bacterial DNA from fecal samples. M.B. performed SFB qPCR. D.A. performed and analyzed behavioral tests. M.B., C.S., D.C.U. and L.S. assisted with animal husbandry, timed pregnancies, and mice genotyping. K.R. advised on behavioral test design and provided equipment for behavioral tests. A.W. provided original mouse strains and funding support. I.A.M. conceptualized and supervised the study. D.A. wrote the manuscript (original draft, review, and editing). A.W. and I.A.M. wrote the manuscript (review and editing). All authors contributed to editing the manuscript.

---

**Supplementary Fig. 1.**
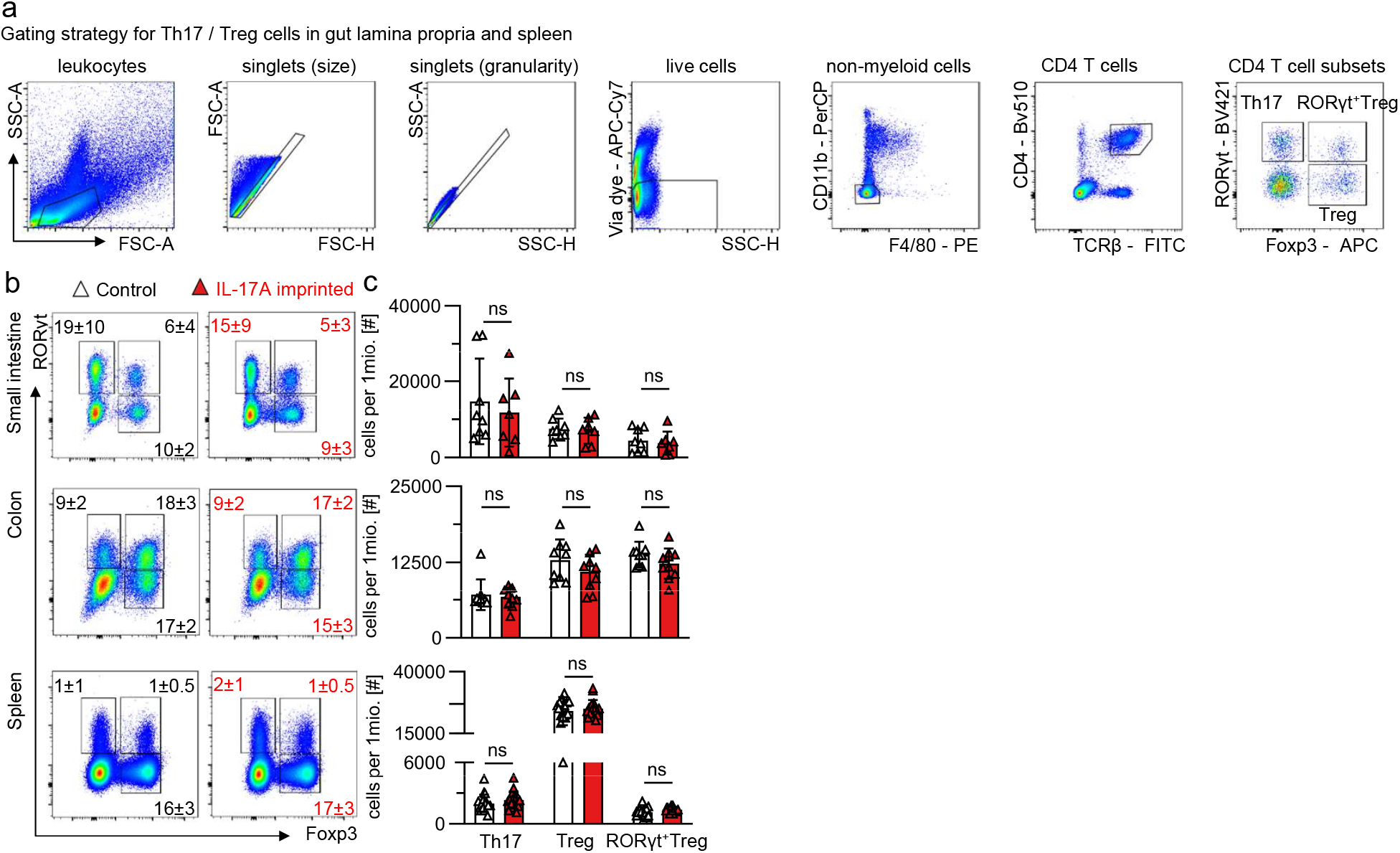
Th17 and Treg cells are not impacted in adult female mice prenatally exposed to IL-17A. **a)** Gating strategy for identification of intestinal lamina propria and splenic Th17/Treg/RORγt+Treg cells by flow cytometry in female mice. **b)** Representative FACS plots depict RORγt and Foxp3 expression by T cells (gated on: CD11b^-^F4/80^-^TCRβ^+^CD4^+^ cells). Numbers within plots indicate mean±SD frequencies of cells in the adjacent gate. No statistically significant differences were noted in frequencies between groups. **c)** numbers of Th17, Treg and RORγt^+^ Treg cells in the small intestine (top), colon (middle) and spleen (bottom). Graphs show pooled data from two (intestines) and four (spleen) independent experiments. Small intestine: n=8 from 2 litters (white) and 7 from 3 litters (red); Colon: n=9 from 4 litters (white) and 10 from 5 litters (red); Spleen: n=17 per group from 6 litters (white) and 8 litters (red). To normalize data from independent experiments, the cell number of each T cell subset was divided by the number of million cells in the sample. Datapoints represent individual mice. Bar graphs indicate mean±SD. ns=p>0.05 two-tailed unpaired t-test. Control=IL-17RA^KO/WT^ offspring born to IL-17RA^KO/KO^ dams (white). IL-17A imprinted=IL-17RA^KO/WT^ offspring born to IL-17RA^KO/KO^IL-17A^OE/WT^ dams (red).

**Supplementary Fig. 2.**
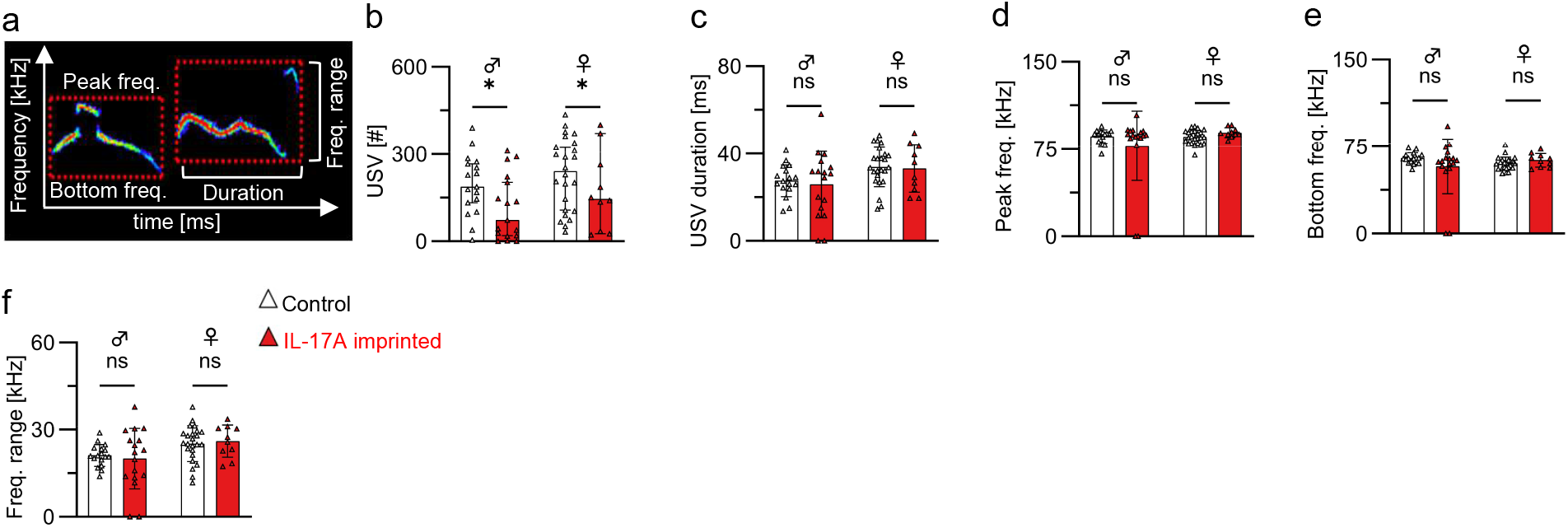
Extended analysis of the ultrasonic vocalization (USV) profile of IL-17A-imprinted pups. a) Example USV profile highlighting the assessed parameters for extended analysis. Red dotted boxes mark individual vocalizations. The quantification shown in c)-e) was performed by averaging the duration, bottom frequency (lower red box edge), and peak frequency (upper red box edge) values of all emitted vocalizations per pup. b) Number of vocalizations. Bar graphs represent median±95%CI. ns=p>0.05 two-tailed unpaired Mann-Whitney test. c) USV duration d) USV low frequency. e) USV high frequency. f) USV frequency range, quantified by subtracting the mean USV low frequency from the mean USV high frequency of an individual pup. b)-f) Data points represent individual pups. Males: n=17 from 6 litters (white) and 17 from 8 litters (red). Females: n=24 from 7 litters (white) and 10 from 8 litters (red). c)-f) Bar graphs indicate mean±SD. *=p<0.05, ns=p>0.05 two-way ANOVA with Šídák’s multiple comparisons test. Control=IL-17RA^KO/WT^ offspring born to IL-17RA^KO/KO^ dams (white). IL-17A imprinted=IL-17RA^KO/WT^ offspring born to IL-17RA^KO/KO^IL-17A^OE/WT^ dams (red).

**Supplementary Fig. 3.**
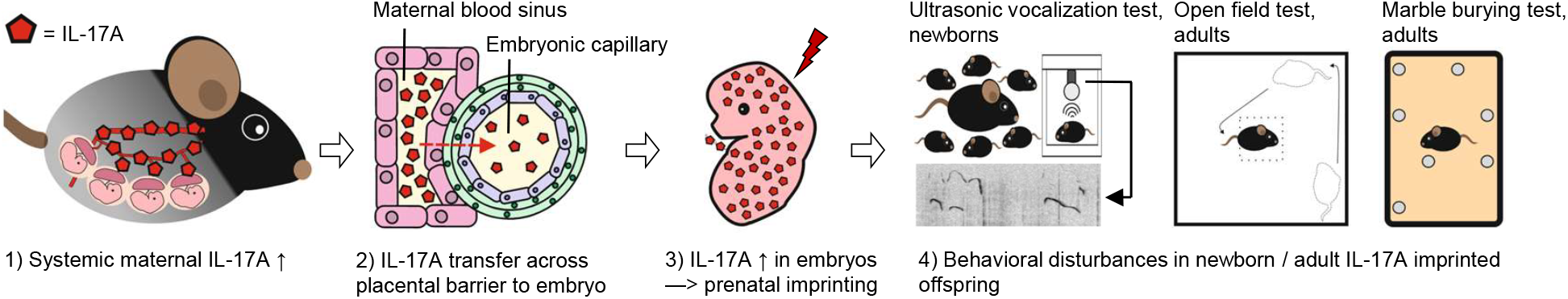
Proposed mechanism by which maternal IL-17A mediates behavioral disturbances in offspring. IL-17A present in the maternal blood (1) is translocated across the placental barrier (2) which is formed by four embryonic cell layers (Pink=Trophoblast giant cell layer; Green=Syncytiotrophoblast bilayer; Blue=Endothelial cell layer). Maternal-derived IL-17A is distributed within the embryo, leading to prenatal imprinting to IL-17A (3). Exposure to IL-17A in the prenatal period leads to long-lasting behavioral disturbances (4).

